# The p150N domain of Chromatin Assembly Factor-1 regulates Ki-67 accumulation on the mitotic perichromosomal layer

**DOI:** 10.1101/082339

**Authors:** Timothy D. Matheson, Paul D. Kaufman

## Abstract

Chromatin Assembly Factor 1 (CAF-1) deposits histones during DNA synthesis. The p150 subunit of human CAF-1 contains an N-terminal domain (p150N) that is dispensable for histone deposition but which promotes the localization of specific loci (Nucleolar-Associated Domains, or “NADs”) and proteins to the nucleolus during interphase. One of the p150N-regulated proteins is proliferation antigen Ki-67, whose depletion also decreases the nucleolar association of NADs. Ki-67 is also a fundamental component of the perichromosomal layer (PCL), a sheath of proteins surrounding condensed chromosomes during mitosis. We show here that a subset of p150 localizes to the PCL during mitosis, and that p150N is required for normal levels of Ki-67 accumulation on the PCL. This activity requires the Sumoylation Interacting Motif (SIM) within p150N, which is also required for the nucleolar localization of NADs and Ki-67 during interphase. In this manner, p150N coordinates both interphase and mitotic nuclear structures via Ki67.

## Introduction

Eukaryotic chromosomes must condense for segregation during mitosis, and then decondense and re-adopt interphase configurations in the next cell cycle.

However, mechanisms that link higher-order mitotic and interphase genome organization remain poorly understood. Notably, cellular heterochromatin is highly enriched at specific sites in interphase nuclei, at the nuclear lamina, in pericentric foci, and in perinucleolar regions (reviewed in (Politz et al., 2016)). Therefore, a major question is how these loci are relocalized after each mitotsis, and what mitotic molecules might aid in this.

Nucleolar Associated Domains (NADs) are genomic loci that interact frequently with the nucleolar periphery (van Koningsbruggen et al., 2010; Matheson and Kaufman, 2015; Németh et al., 2010; Padeken and Heun, 2014). NADs are highly enriched for repetitive DNA satellites and histone modifications associated with heterochromatic silencing such as H3K27me3, H3K9me3, and H4K20me3 (Németh et al. 2010; Politz et al. 2016). One protein recently shown to regulate NAD localization is the p150 subunit of Chromatin Assembly Factor 1 (CAF-1) (Smith et al., 2014). CAF-1 is a highly conserved three-subunit protein complex which deposits histone (H3/H4)_2_ tetramers onto replicating DNA during S-phase of the cell cycle (Kaufman et al., 1995; Krude, 1995; Smith & Stillman, 1989), and is particularly important for DNA replication and maintenance of histone marks at heterochromatic loci (Baldeyron et al., 2011; Dohke et al., 2008; Houlard et al., 2006; Huang et al., 2010; Quivy et al., 2004, 2008; Reese et al., 2003; Sarraf and Stancheva, 2004). In addition to these functions, the N-terminal domain of p150 (p150N) regulates the association of NADs to the perinucleolar region, and also regulates the nucleolar localization of several proteins including the proliferation antigen Ki-67 (Smith et al., 2014).

Ki-67 also regulates heterochromatin modification (Sobecki et al., 2016) and affects its condensation (Kametaka et al., 2002; Takagi et al., 1999). Ki-67 is highly expressed in proliferating cells, minimally expressed in quiescent cells (Gerdes et al., 1984; Gerdes et al., 1983), and meta-analyses of numerous clinical studies have validated Ki-67 as a prognostic marker in grading tumors (Luo et al., 2015; Pezzilli et al., 2016; Pyo et al., 2016; Richards-Taylor et al., 2016). Similar to Ki-67, p150 is also highly expressed in proliferating cells, minimally expressed in quiescent cells, and has been proposed as an alternative cellular proliferation marker in clinical studies (Polo et al., 2004).

Recent studies have demonstrated that Ki-67 is required for formation of the perichromosomal layer (PCL) (Booth et al., 2014; Sobecki et al., 2016), a layer of proteins that surrounds all condensed chromosomes during mitosis (reviewed in (Van Hooser 2005)). At the PCL, Ki-67’s status as a large, charged protein that possesses “surfactant”-like properties keeps individual mitotic chromosomes dispersed rather than aggregated upon nuclear envelope disassembly, thereby ensuring normal kinetics of anaphase progression (Cuylen et al., 2016). Thus, Ki-67 is important for mitotic chromosome structure and function. Because p150N regulates interphase Ki-67 localization, we investigated here whether p150N also affects Ki-67 localization during mitosis.

## Results and Discussion

### Ki-67 regulates alpha satellite localization to the perinucleolar region

Recent studies have suggested Ki-67 regulates repetitive DNA localization to the nucleolus. Ki-67 depletion decreased the nucleolar association of a LacO array proximal to the rDNA repeats on chromosome 13 (Booth et al., 2014), and the centromeric histone variant CENP-A (Sobecki et al., 2016). To determine if Ki-67 also regulates satellite repeat association with the nucleolus, we transfected siRNAs into HeLa cells and performed immuno-FISH to visualize the position of alpha satellite DNA from chromosome 17 (αSat 17) relative to the nucleolar protein fibrillarin. HeLa cells transfected with siRNAs directed against luciferase or a scramble control displayed ~45% of αSat 17 alleles associated with nucleoli, which represents a high-frequency interaction characteristic of a NAD locus (Németh et al., 2010; Smith et al., 2014; van Koningsbruggen et al., 2010). In contrast, cells transfected with two different siRNA reagents directed against distinct regions of the Ki-67 mRNA displayed αSat 17 association frequencies to ~25% (Figure 1, A and B).

**Figure 1.**
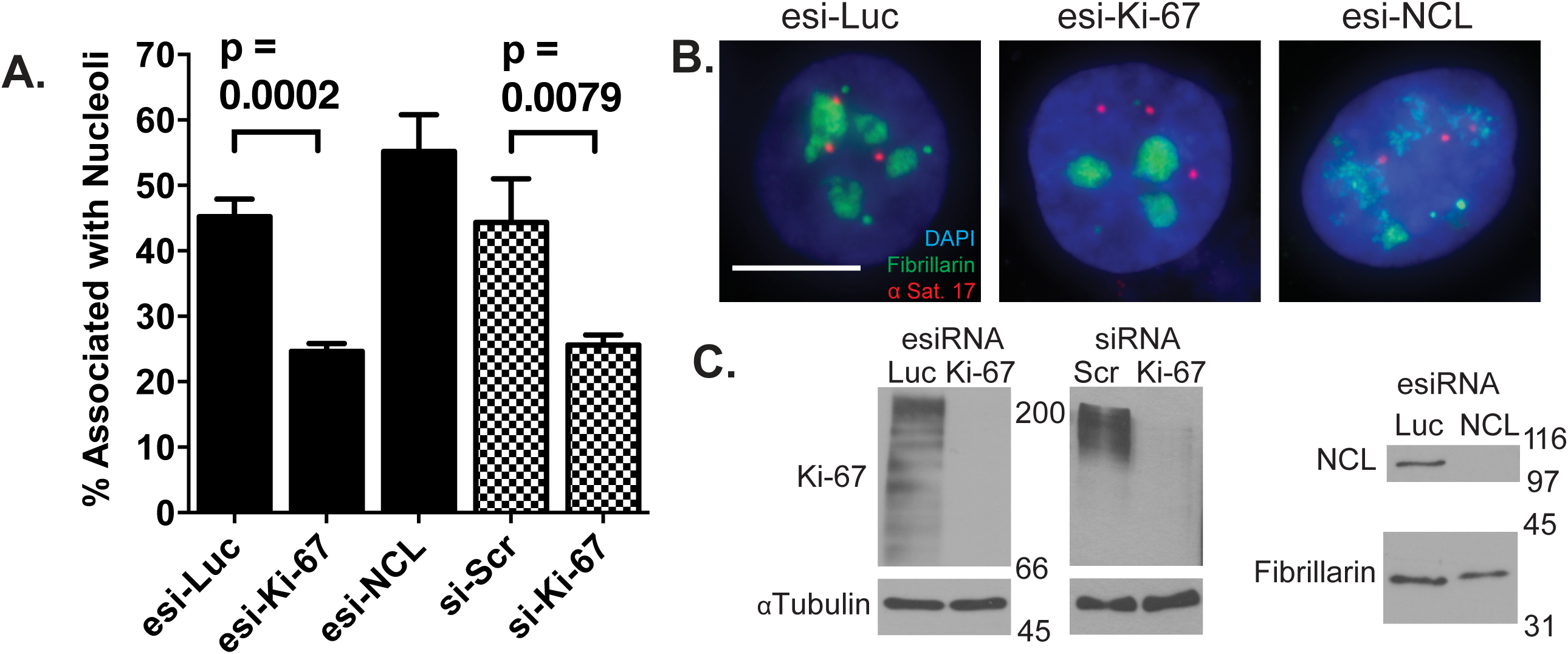
Alpha-satellite associations with nucleoli are reduced upon depletion of Ki-67 but not nucleolin. (A): Immuno-FISH analyses of the association between alpha satellite DNA from chromosome 17 (αSat 17, red) and fibrillarin (green) was performed in HeLa cells transfected for 72 hours with in vitro-diced “esiRNAs” targeting luciferase, Ki-67, or nucleolin (NCL) (black bars), or with synthetic duplex siRNAs (checked bars) targeting Ki-67 or a scrambled sequence control (si-Scr). The percentage of αSat 17 alleles co-localized with fibrillarin is indicated, with means and standard deviation error bars from three replicate experiments. p values comparing association frequencies in cells treated with esi-Luc (N=669 alleles) vs. esi-Ki-67 (573 alleles), and si-SCR (N=540 alleles) with si-Ki-67 (N=567 alleles) are indicated. (B): Representative FISH images of cells analyzed in panel (A). Scale bar is 10 µm. (C): Immunoblot analyses of cells from panel (A). Blots were probed with antibodies recognizing Ki-67, α-Tubulin (loading control), NCL, or fibrillarin (loading control), as indicated.

We also depleted HeLa cells of nucleolin (NCL), a nucleolar protein which primarily localizes to the perinucleolar region (Bugler et al., 1982) and regulates transcription, folding, and assembly of ribosomal RNA (Ginisty et al., 1999; Ginisty et al., 1998; Rickards et al., 2007; Roger et al., 2002). Depletion of NCL altered nucleolar morphology (Figure 1, B) as previously reported (Ma et al., 2007; Ugrinova et al., 2007) but did not decrease the association of αSat 17 with fibrillarin (Figure 1, A). We conclude that Ki-67 but not all nucleolar proteins are required for efficient localization of NADs to nucleoli.

### A subset of p150 co-localizes with Ki-67 foci during mitosis and early G1 phases

Ki-67 has three distinct nuclear localization patterns, which appear at distinct cell-cycle periods. First, Ki-67 localizes to the nucleolus during interphase (Cheutin et al., 2003; Traut et al., 2002; Verheijen et al., 1989), and a subset co-localizes with p150 at the perinucleolar region (Smith et al., 2014).

Second, Ki-67 localizes to the PCL during mitosis (Gerdes et al., 1984; Gerdes et al., 1983; Verheijen, et al., 1989). In contrast to the chromosome-associated state of mitotic Ki-67, mitotic phosphorylation evicts the majority of CAF-1 from mitotic chromosomes and inhibits its nucleosome assembly activity (Keller and Krude, 2000; Marheineke and Krude, 1998; Matsumoto-Taniura et al., 1996; Murzina et al., 1999). However, mass spectrometric analyses suggested that a mitotic chromosome-associated subset of p150 exists (Ohta et al. 2010), but its relationship to the PCL had not been directly tested. To analyze this we took advantage of the fact that Ki-67 localization to the PCL remains visibly unchanged even after extraction with 2 M NaCl and treatment with either RNase A or DNase I (Sheval and Polyakov, 2008), indicating the structural stability of the PCL is independent of the chromatin it surrounds. We performed immunofluorescence on RPE1-hTERT cells that were digested with either RNase A, DNase I, or mock digested, and then high-salt extracted. As previously reported, Ki-67 remained on the PCL throughout mitosis even after DNA or RNA digestion, as shown for prophase (Figure 2, A), metaphase (Figure 2, B), and anaphase (Figure 2, C). As expected, p150 as well as Ki67 could be easily detected on anaphase chromosomes presumably because at that stage the mitotic phosphorylations of CAF-1 are removed, triggering bulk re-association with chromatin and reactivation of nucleosome assembly activity (Keller and Krude, 2000). In contrast, during prophase or metaphase a large amount of p150 evicted into the nucleoplasm made it difficult to determine whether p150 localized to the PCL. The combination of high-salt extraction and DNase I digestion however revealed that a subset of p150 localized to the PCL during all phases of mitosis.

**Figure 2.**
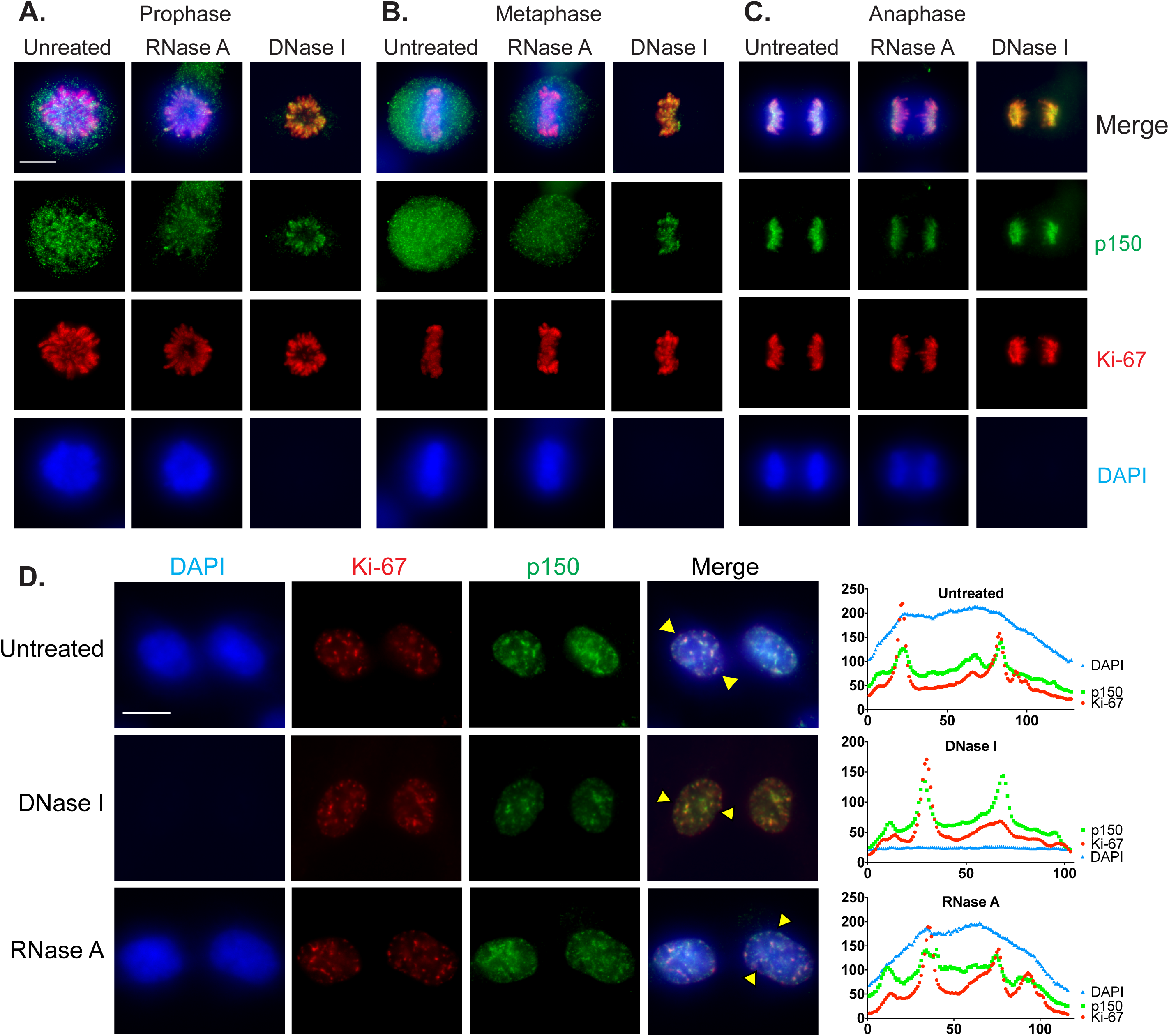
A subset of p150 localizes to the PCL during mitosis and to Ki-67 foci during early G1 of interphase. RPE1-hTERT cells were either untreated or digested with RNase A or DNase I as indicated, and then high-salt extracted. Cells were stained with DAPI to detect DNA (blue), and with antibodies recognizing p150 (green), and Ki-67 (red). Cells from prophase (A), metaphase (B), and anaphase (C) are shown. Note the DNase I-treated cells lack DAPI staining. Scale bar is 10 µm. (D) p150 co-localizes with Ki-67 foci during early G1. RPE1-hTERT cells were treated and stained as indicated above. Pairs of recently divided cells featuring hundreds of Ki-67 foci characteristic of early G1 are shown here. Line scans (right-hand panels) of individual cells (yellow triangles in merged images) were used to assess co-localization of p150 with Ki-67 signal. Scale bar is 10 µm.

The third Ki-67 localization pattern occurs after cytokinesis and early in G1 phase. At this stage, Ki-67 localizes to hundreds of distinct foci (du Manoir et al., 1991; Isola et al., 1990) which co-localize with heterochromatic satellite repeats (Bridger et al., 1998). These foci are transient and appear to be intermediates in the process of reformation of nucleoli as cells transition from mitosis to interphase (Saiwaki et al., 2005). Because a subset of p150 is in the mitotic PCL (Figure 2, A-C), we sought to determine if p150 also co-localized with early G1 Ki-67 foci. To do this, we performed immunofluorescence on early G1 RPE1-hTERT cells digested with nucleases or mock digested as previously described (Figure 2, D). In all three conditions, a subset of p150 foci in early G1 cells co-localized with Ki-67 foci, as indicated by line scan analyses.

These data suggest that a subset of p150 travels together with Ki67 during exit from mitosis, as PCL components transit towards reforming nucleoli.

### p150 regulates Ki-67 accumulation on the Perichromosomal Layer (PCL)

Our previous work demonstrated that p150 is required for normal steady-state accumulation of Ki-67 in the nucleolus during interphase (Smith et al., 2014). Because Ki-67 is essential for the formation of the perichromsomal layer (PCL) in mitotic cells (Booth et al., 2014; Sobecki et al., 2016), we tested whether p150 also regulated Ki-67 localization during mitosis. Via immunofluorescence, we examined Ki-67 distribution in HeLa S3 cells expressing an inducible shRNA directed against either luciferase (Luc) or p150 (Figure 3, A). As expected, we found Ki-67 robustly associated with the PCL during all phases of mitosis in the negative control cells expressing sh-Luc. In contrast, cells expressing sh-p150 displayed less Ki-67 staining on the PCL, as demonstrated in the exposure times matched with the sh-Luc samples. When exposure times were increased, Ki-67 was detected on the PCL, indicating that the PCL was not entirely disassembled upon p150 depletion. When the Ki-67 fluorescence was quantified in prometaphase cells from three biological replicates, cells expressing sh-p150 displayed, on average, a 3.5-fold decrease in fluorescence intensity compared to the control cells expressing sh-Luc (Figure 3, B). To test whether this effect of p150 depletion could result from global masking of epitopes on the PCL, we co-stained some samples with an antibody recognizing the mitotic histone modification H3-S28ph (Goto et al., 1999; Goto et al., 2002). As shown in the prometaphase cell in Figure 3A, this antibody stained cells expressing sh-Luc or sh-p150 equally well, suggesting that p150 regulates Ki-67 accumulation on the PCL rather than affecting overall epitope accessibility on mitotic chromosomes.

**Figure 3.**
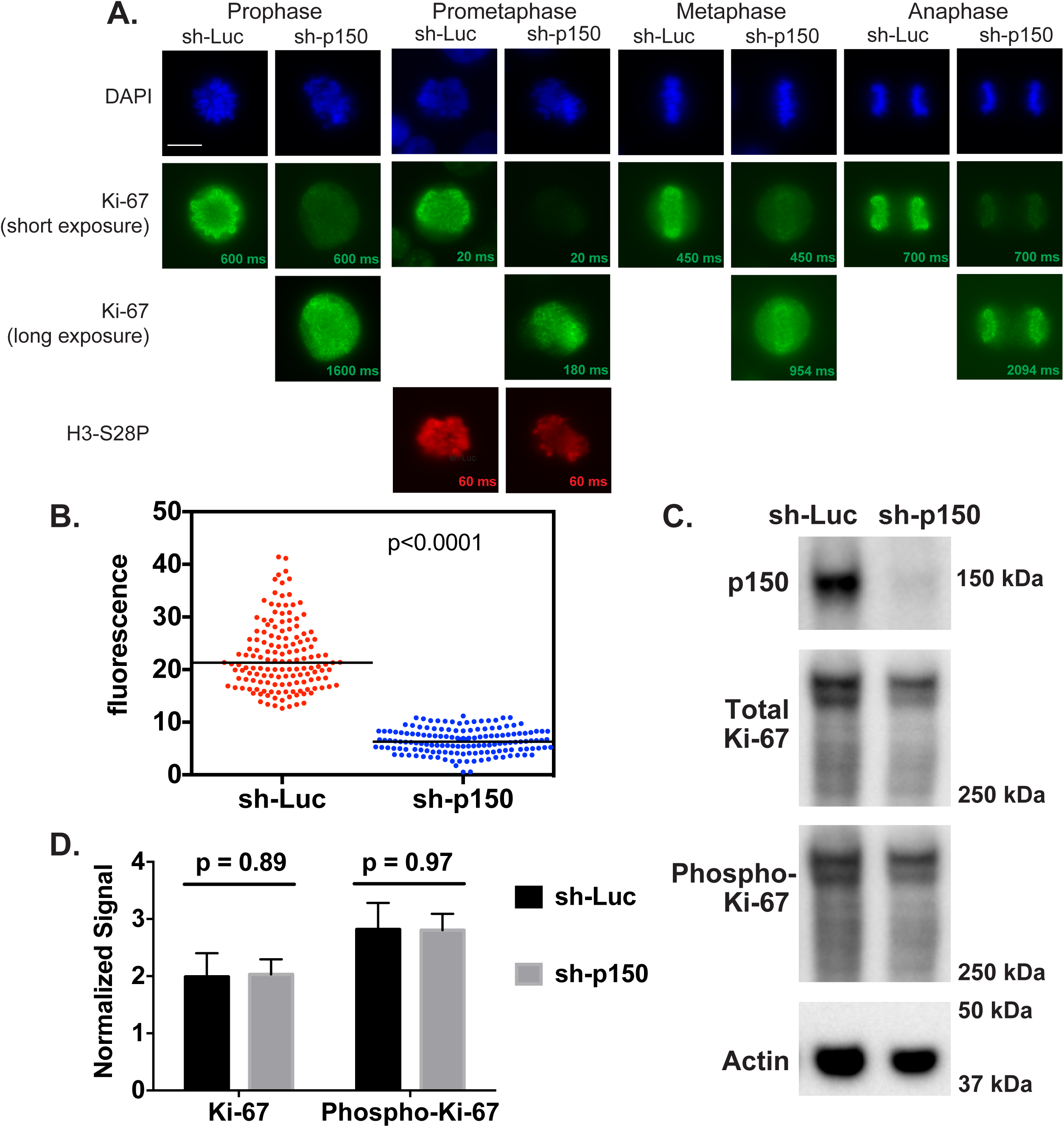
p150 regulates Ki-67 localization to the PCL during mitosis. (A): HeLa S3 cells from the indicated cell cycle stages expressed an shRNA directed against luciferase (sh-Luc) or p150 (sh-p150) for 72 hours and were stained with DAPI to visualize DNA (blue) and antibodies against Ki-67 (green). Exposure times for Ki67 are indicated on each image, and in the sh-p150 expressing cells different exposure times are shown to illustrate reduced Ki-67 accumulation on the PCL. As a positive control for antibody accessibility, cells in prometaphase were also stained with antibodies recognizing the mitotic marker histone H3-S28-phosphate (red). Scale bar is 10 µm. (B): Quantified corrected total cellular fluorescence of cells from three biological replicate experiments of cells expressing sh-luc (red, N=150) or sh-p150 (blue, N=147). (C): Immunoblot analysis of extracts from shRNA-expressing cells described in (A) arrested in mitosis (12 hours in 100 ng/mL nocodazole followed by shaking off mitotic cells). Blots were probed with antibodies recognizing p150 (top), Ki-67 (upper middle), phospho-Ki-67 (lower middle), and actin (loading control, bottom). Numbers on the right indicate migration of marker proteins, in kDa. (D): Quantification of the Ki-67 and phospho-Ki-67 blots from (C), normalized to actin signal (N=3). Quantification was performed using the BioRad ChemiDoc system.

To explore how p150 may be regulating Ki-67 localization, we tested whether p150 is required for maintaining steady-state levels of Ki-67. Our previous work demonstrated that Ki-67 steady-state levels were not affected upon p150 depletion in asynchronous cells (Smith et al., 2014). To distinguish whether p150 regulates the steady-state protein levels of Ki-67 during mitosis, extracts from mitotic-arrested cells expressing sh-Luc or sh-p150 were collected and analyzed by immunoblotting (Smith et al., 2014) (Figure 3, C and D). Total levels of Ki-67 in mitotic samples normalized to actin signals were not significantly changed upon p150 depletion (Figure 3, C and D), indicating that p150 is not required for maintaining steady-state levels of Ki-67 during mitosis. At the beginning of mitosis, Ki-67 localization to the PCL occurs in conjunction with hyperphosphorylation of the Ki-67 protein (Endl and Gerdes, 2000; MacCallum and Hall, 1999; Takagi et al., 2014). These phosphorylations are important for PCL localization, as treatment of mitotic cells with protein kinase inhibitors results in dephosphorylation of Ki-67 and relocalization of Ki-67 away from the PCL to distinct nuclear foci (MacCallum and Hall, 1999). Therefore, we tested whether p150 regulates Ki-67’s mitotic phosphorylation status by reprobing the immunoblots of mitotically-arrested cell extracts with antibodies that specifically recognize Ki-67 phosphorylated on Cdk consensus sites within the Ki-67 internal repeat structure (Takagi et al., 2014).

However, we detected no statistically significant changes upon p150 depletion (Figure 3, C and D). Therefore, p150 does not appear to regulate the steady-state levels or phosphorylation of Ki-67 during mitosis.

### The SIM within the N-terminus of p150 is required for Ki-67 localization to the PCL

To map p150 domains required for regulating Ki-67 accumulation on the PCL, we utilized previously published HeLa cell lines stably expressing V5-epitope tagged, sh-RNA resistant, p150-derived transgenes (Smith et al., 2014). These V5-transgene cell lines were acutely depleted of endogenously-encoded p150 via lentiviral expression of sh-p150 for 72 hours prior to immunofluorescence analysis of Ki-67.

The C-terminal two-thirds of p150 serves as the scaffold for binding the other two subunits of the CAF-1 complex, and thereby is essential for CAF-1’s nucleosome assembly activity (Kaufman et al., 1995; Takami et al., 2007). In contrast, p150’s N-terminus is dispensable for nucleosome assembly (Kaufman et al., 1995) but is necessary and sufficient to maintain Ki-67 localization to the nucleolus during interphase (Smith et al., 2014). We found that depletion of p150 in HeLa cells expressing either Luciferase or a p150 transgene encoding the C-terminus (amino acids 311-938) displayed reduced Ki-67 localization to the PCL (Figure 4, A). In contrast, cells expressing either full-length p150 (aa 1-938) or only the N-terminus (p150N, aa 1-310) maintained normal amounts of Ki-67 on the PCL (Figure 4, A). Therefore, the p150 N-terminus regulates mitotic Ki-67 abundance in a manner independent of the chromatin assembly activity of the CAF-1 complex.

**Figure 4.**
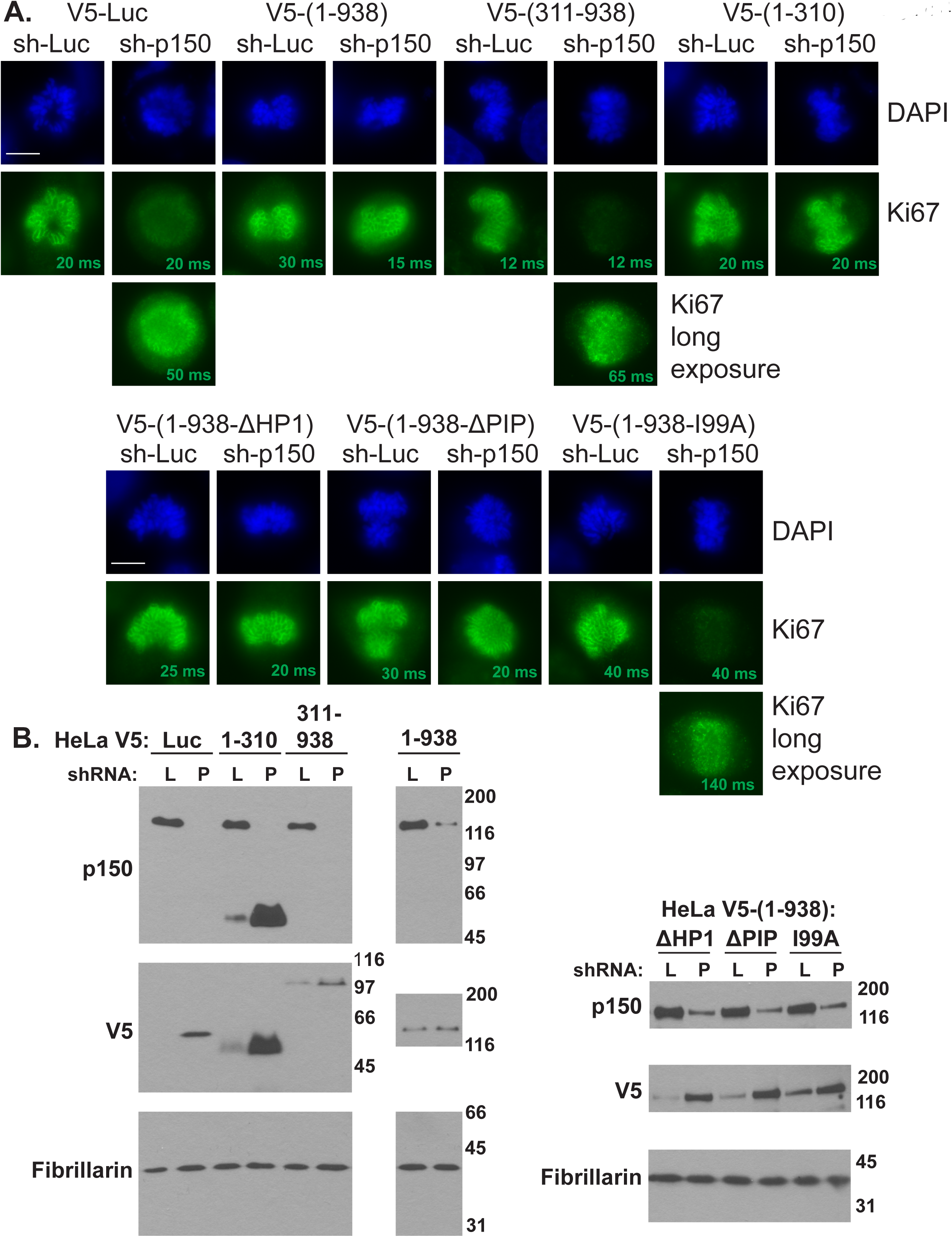
The SIM within the N-terminal domain of p150 is sufficient to maintain Ki-67 localization to the PCL. (A): HeLa cells in prometaphase expressing the indicated shRNA-resistant p150 transgene were infected with lentiviruses encoding the indicated shRNAs (sh-Luc or sh-p150) for 72 hours, and then stained with DAPI to visualize DNA (blue) and antibodies against Ki-67 (green). Note that upon expression of sh-p150, cells expressing luciferase (V5-Luc), the p150 C-terminus (V5-(p150-311-938)), or p150 with a I99A point mutation in the SIM motif required longer exposure times to detect Ki-67 on the PCL. Scale bar is 10 µm. (B): Immunoblots of cell extracts described in (A), expressing either sh-Luc (L) or ph-p150 (P). Blots were probed with antibodies recognizing p150 (top), V5 transgenes (middle), and fibrillarin (loading control, bottom). Note the depletion of endogenous p150 in sh-p150 lanes, and that V5-tagged transgene expression often increased in these lanes, as we had observed previously (Smith et al., 2014).

p150N contains several known interaction motifs, including a non-canonical PCNA-interacting peptide (PIP) (Moggs et al., 2000; Rolef Ben-Shahar et al., 2009), a heterochromatin protein 1-binding domain (Murzina et al., 1999), and a sumoylation-interacting motif (SIM) (Sun and Hunter, 2012; Uwada et al., 2010). Notably, the SIM in p150N is required for normal Ki-67 localization to the nucleolus during interphase (Smith et al., 2014). We therefore determined if transgenes encoding full-length p150 with mutations in the three motifs described above supported normal Ki-67 accumulation during mitosis. The cell lines expressing p150 transgenes with mutations in the PIP and HP1 domains maintained normal Ki-67 levels on the PCL. However, cells expressing a p150 transgene with a single amino acid mutation (I99A) that disrupts the SUMO-binding activity of the SIM (Uwada et al., 2010) displayed reduced levels of Ki-67 on the PCL (Figure 4, A). Together, our data indicate that that the SIM motif within the N-terminus of p150 is required for the normal accumulation of Ki67 on the PCL during mitosis.

This study therefore suggests a hierarchical relationship between p150N and Ki-67 in regulating nuclear architecture during both interphase and mitosis. p150 and Ki-67 both co-localize during mitosis and interphase, regulate NAD localization, and have established roles in regulating heterochromatin. p150N appears to function upstream of Ki-67 in this hierarchy, as p150N is required for efficient Ki-67 association with the PCL during mitosis (Fig. 4) and with the nucleolus during interphase (Smith et al., 2014). p150N is dispensable for nucleosome assembly by CAF-1, indicating that p150’s role in regulating Ki-67 localization is independent of CAF-1 activity. During both mitosis and interphase, the SIM within p150N is required for Ki-67 localization, suggesting that this action involves an as yet unidentified sumoylated protein. These data suggest that future studies should explore the contributions of the p150N domain and its putative sumoylated binding partners to the relationship between Ki-67 and p150 in regulating mitotic and interphase nuclear structure.

## Materials and Methods

### Cell culture

HeLa S3 cells containing the Trex CMV/TO promoters driving either sh-Luc or sh-p150 expression (Campeau et al., 2009) were maintained in RPMI media with 5% tetracycline-free fetal bovine serum (FBS), 2 mM L-glutamine, and antibiotic/antimycotic solution (Life Technologies). shRNA expression was induced by supplementing media with 2 µg/mL doxycycline for 72 hours prior to fixation or processing. Mitotic cells were enriched by adding 100 ng/mL nocodazole for 12 hours (hours 60-72 of the doxycycline treatment) and then freed from the cell culture flask surface by vigorous shaking. HeLa cells continuously expressing the V5-tagged p150 transgenes (Smith et al., 2014) were maintained in DMEM and supplemented with 10% FBS, 2 mM L-glutamine, and antibiotic/antimycotic solution (Life Technologies). RPE1-hTERT cells (a generous gift from Judith Sharp) were maintained in 50:50 DMEM-F12 media supplemented with 10% FBS, 7.5% sodium bicarbonate, 2 mM L-glutamine, and antibiotic/antimycotic solution (Life Technologies).

### Depletion Reagents

For lentiviral depletion (Smith et al., 2014), cells were infected at MOI = 7.5 with 6 µg /mL polybrene for 72 hours. Lentivirus reagents were synthesized as previously described (Campeau et al., 2009; Smith et al., 2014). esiRNA reagents were generated as previously described (Fazzio et al., 2008; Smith et al., 2014), using primers listed in Table 2 to generate dsRNA. For esiRNA transfection, 500 ng of siRNAs were transfected in 1 ml Opti-MEM (Life Technologies) with 6 µl of Oligofectamine (Life Technologies). After 6.5 h, 2.5 ml of media was added on top of the transfection cocktail and cells were processed after 72 hours. For synthetic siRNA transfection, 10 µL of 5 µM scramble (Ambion, AM4611) or Ki-67 (Ambion, Catalog #4392420-s8796) siRNAs were diluted in 400 µL Opti-MEM (Life Technologies) with 5 µL RNAiMAX (Invitrogen) and incubated at room temperature for 10 minutes. The siRNA cocktail was then slowly added to 6-well dishes containing 800 µL Opti-MEM, and 6 hours later 2.5 mL of appropriate media (lacking antibiotics) was added. Note that the esiRNA directed against Ki-67 targets the Ki-67 repeat region within exon 13 while the synthetic siRNA targets nucleotides 559-577 (CGUCGUGUCUCAAGAUCUAtt) within exon 6.

**Table 1:**
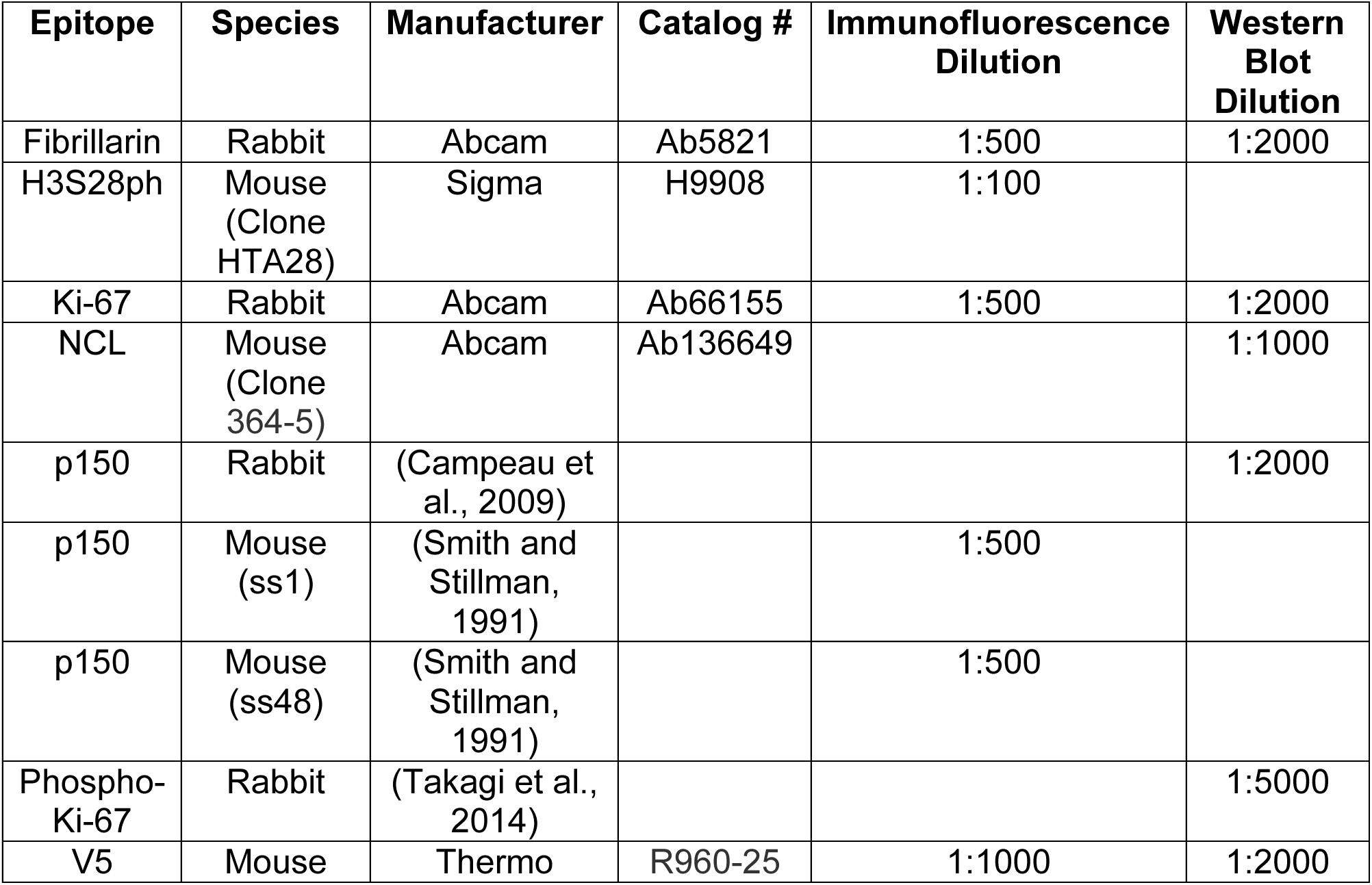
Antibodies.

**Table 2:**
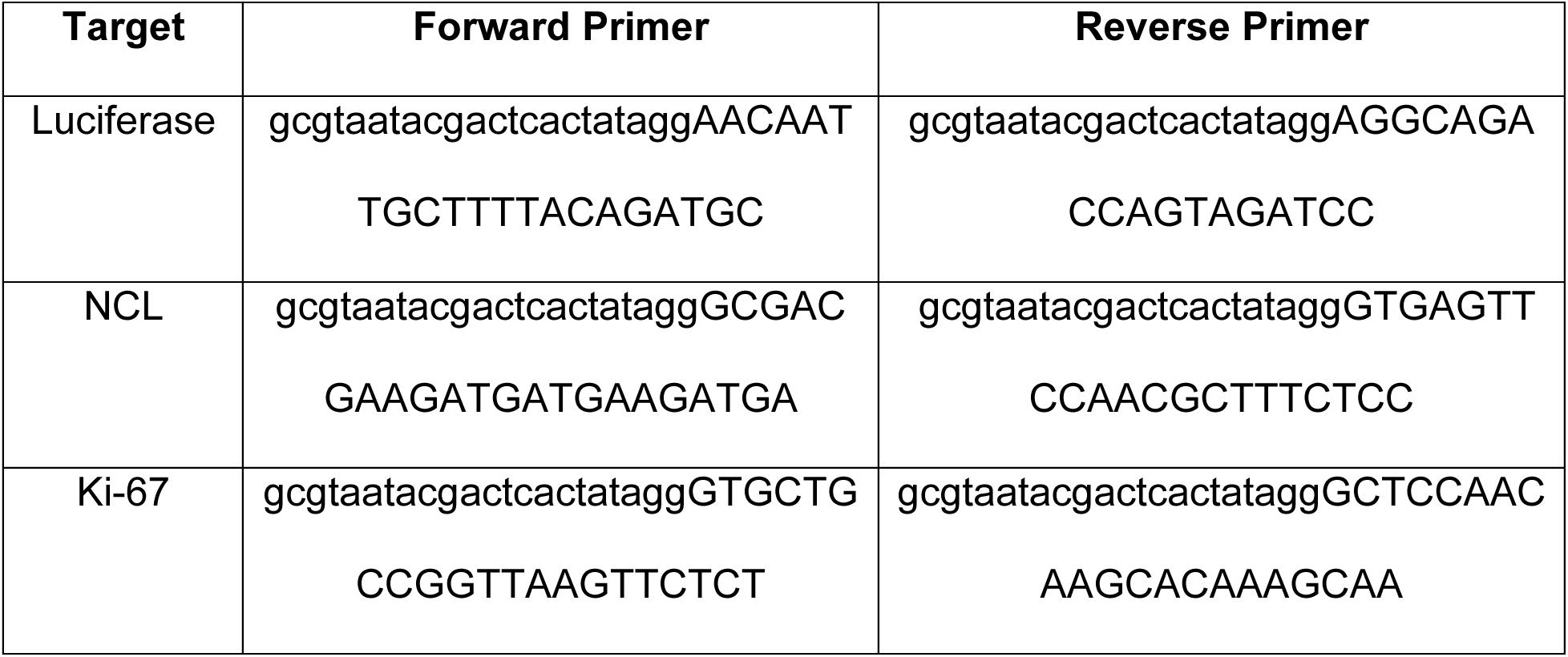
esiRNAs.

### Immunofluorescence and Immuno-FISH

Cells were plated on poly-lysine (Sigma) coated coverslips for 24 hours prior to manipulation. For immunofluorescence experiments, fixation and permeabilization was performed in 6-well tissue culture dishes to minimize displacement of mitotic cells. After aspiration of media, cells were immediately fixed with 4% paraformaldehyde/PBS for 10 minutes at room temperature. Cells were then gently washed with ice-cold PBS and permeabilized with 0.5% Triton TX-100/PBS on ice for 5 minutes. Cells were then washed twice with room temperature PBS and then incubated in blocking buffer (1% BSA/PBS) for 5-30 minutes in a 37 °C humid chamber. After blocking, cells were incubated with primary antibody in blocking buffer for 2 hours in a 37 °C humid chamber. Coverslips were then transferred to Coplin jars and washed three times with PBS on a rotating platform for 10 minutes at room temperature. After washing, cells were incubated with secondary antibody in blocking buffer for 1 hour in a 37°C humid chamber. Coverslips were then transferred to Coplin jars and washed three times with PBS on a rotating platform for 10 minutes at room temperature. After the final washing step, cells were transferred to a Coplin jar containing DAPI (50 ng/mL) in PBS for 2 minutes. Coverslips were mounted with Vecta Shield (Vector Labs) and images were taken on a Zeiss Axioplan2 microscope with a 63x objective. Corrected Total Cell Fluorescence (CTCF) was quantified using the Image J “measure” feature and background subtracted from an area of the image without a cell. Images were all captured using the same exposure time across 3 biological replicates. Immuno-FISH was performed as previously reported (Smith et al., 2014).

### Nuclease Digestion

RPE1-hTERT cells were utilized for these experiments rather than HeLa cells because they are sufficiently adherent during detergent permeabilization/ nuclease digestion/ high salt extraction. Nuclease digestion was performed as previously reported (Sheval and Polyakov, 2008), with some modification. Cells were plated onto poly-lysine coated coverslips in 6-well dishes at least 24 hours before manipulation, and all subsequent steps were performed with solutions containing 100 µM PMSF. Cells were washed once with Digestion buffer (10 mM Tris-HCl, pH 7.6, 5 mM MgCl_2_, 1 mM CuSO_4_), then incubated in 1% Triton TX-100/ Digestion buffer for 10 minutes at 4 °C. Cells were then gently washed with digestion buffer and incubated in 100 µg/ml DNase I/ Digestion buffer, 100 µg/ml RNAse A/ Digestion buffer, or mock digested (“untreated”) in Digestion buffer in a 37°C humid chamber for 30 minutes. Proteins were then extracted by incubating for 10 minutes in Extraction buffer (2M NaCl, 10 mM EDTA, 20 mM Tris-HCl, pH 7.6) at 4°C. Cells on coverslips were immediately fixed as described above, or RNA samples were processed using Trizol (Invitrogen). Briefly, 1 mL Trizol was added directly to the plate and cells were homogenized by pipetting. After incubating for 5 minutes at room temperature, samples were stored at −80 °C until further processing.

After thawing, 200 µL Chloroform was added and samples were shaken vigorously for 15 seconds before incubation at room temperature for 2 minutes. Samples were then centrifuged at 13000 x g for 15 minutes at 4 °C. The aqueous phase was transferred to a fresh tube and 1 volume of isopropanol was added and mixed well by vigorous shaking. Samples were then incubated at room temperature for 10 minutes before centrifugation at 13000 x g for 15 minutes at 4 °C. Samples were then washed with 75% ethanol and resuspended with nuclease-free water. 10% of recovered samples were run on a 1% agarose gel to compare RNA digestion efficiency.

Early G1 cells were selected by choosing pairs of cells which appeared significantly smaller than surrounding cells and featured hundreds of Ki-67 foci (Croft et al., 1999; Isola et al., 1990; du Manoir et al., 1991). Line scans were performed using the RGB Profiles Tool plugin for Image J.

### Western blotting

15 µg of protein were loaded and run through a either Tris-HCl polyacrylamide gradient gel (20-5%) for asynchronous samples, or through a NuPAGE™ Novex™ 3-8% Tris-Acetate Protein Gel (Thermo Fischer) for mitotically-arrested samples. Protein samples were then transferred to a PVDF membrane at 20 volts for 17 hours at 4°C in order to maximize the transfer of high molecular weight Ki-67. Chemiluminescence was acquired using the Biorad ChemiDoc system, and quantified using Biorad Image Lab V 5.2.1.

## Acknowledgements

We would like to thank Michael Brodsky for the generous use of the AxioPlan microscope, Masatoshi Takagi and Naoko Imamoto for the anti-phospho-Ki-67 antibody, and Tom Fazzio for fruitful discussions. This work was supported by NIH grants R01 GM55712 and U01 DA040588 to PDK.

---

**Supplemental Figure 1.**
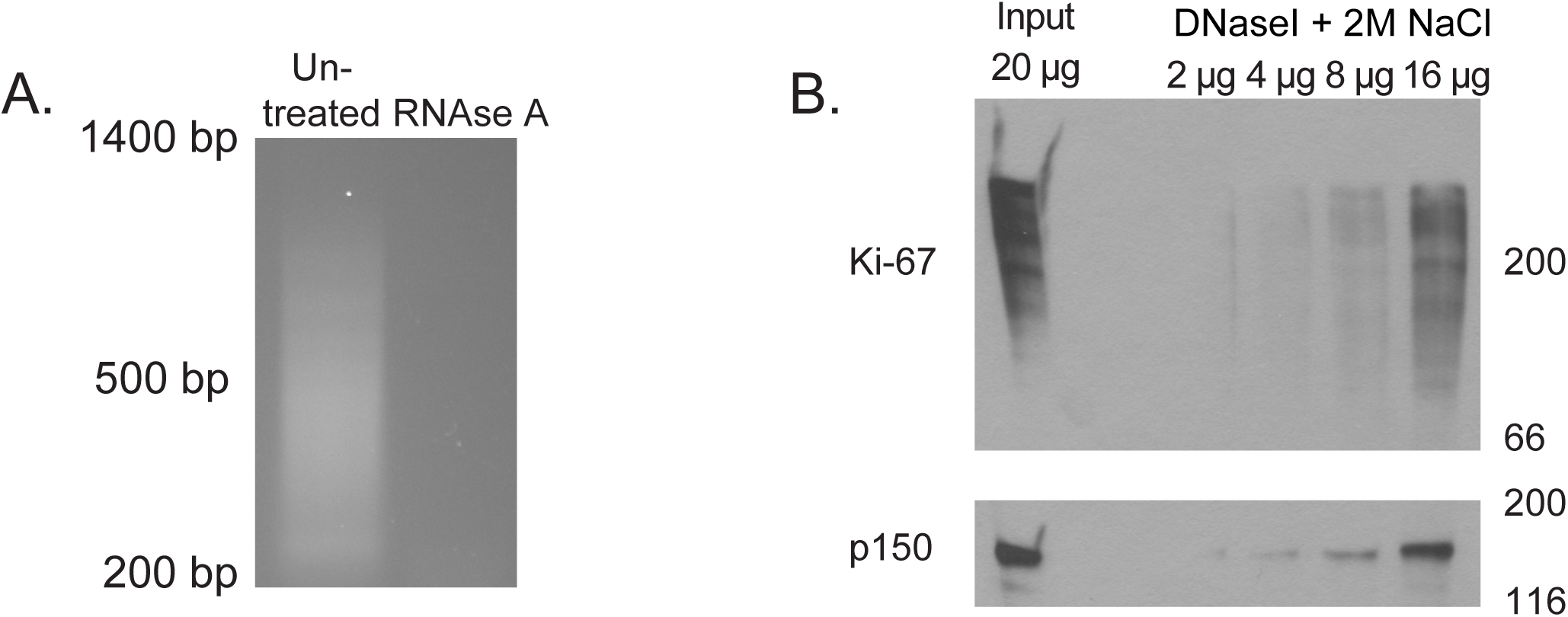
RNase A mediated degradation and western blot of DNase I treated PCL preparation. (A): Positive control for RNase A treatment. RNA from untreated and RNase A-treated cells analyzed in Figure 2 was purified using Trizol reagent and analyzed on a 1% agarose gel. (B): Western blot of mitotic HeLa S3 cells (Input lane, 20 µg) or mitotic HeLa S3 cells digested with DNase I and salt extracted as in Figure 2 (2 µg, 4 µg, 8 µg, 16 µg as indicated). Blot was probed with antibodies recognizing Ki-67 (top) and p150 (bottom). Numbers on the right indicate migration of marker proteins, in kDa.

**Supplemental Figure 2.**
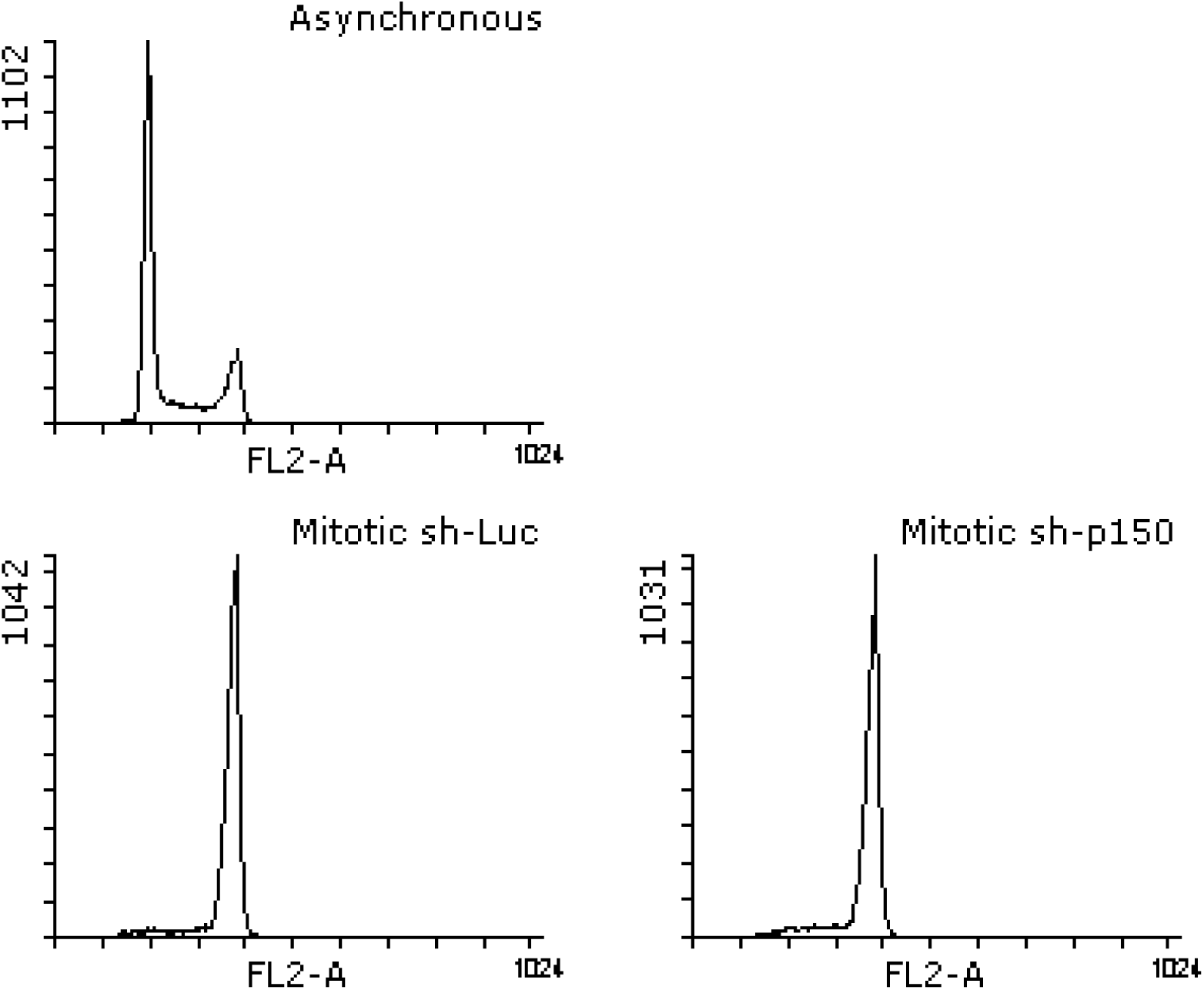
FACS histograms of asynchronous and mitotic-arrested cells. FACS histograms showing cell-cycle profiles of propdium-iodide stained HeLa S3 samples from Figure 3. Note the enrichment of the G2/M peak in mitotic samples (sh-Luc, bottom left; sh-p150, bottom right) compared to the asynchronous sample (top panel). Mitotic cells were enriched by treating with 100 ng/mL nocodazole for 12 hours prior to vigorously shaking cell culture flasks and collecting media.

## References

Baldeyron, C., Soria, G., Roche, D., Cook, A.J.L., and Almouzni, G. (2011). HP1alpha recruitment to DNA damage by p150CAF-1 promotes homologous recombination repair. J. Cell Biol. 193, 81–95.

Booth, D.G., Takagi, M., Sanchez-Pulido, L., Petfalski, E., Vargiu, G., Samejima, K., Imamoto, N., Ponting, C.P., Tollervey, D., Earnshaw, W.C., et al. (2014). Ki-67 is a PP1-interacting protein that organises the mitotic chromosome periphery. Elife 3, e01641.

Bridger, J.M., Kill, I.R., and Lichter, P. (1998). Association of pKi-67 with satellite DNA of the human genome in early G1 cells. Chromosom. Res. 6, 13–24.

Bugler, B., Caizergues-Ferrer, M., Bouche, G., Bourbon, H., and Amalric, F. (1982). Detection and localization of a class of proteins immunologically related to a 100-kDa nucleolar protein. Eur. J. Biochem. 128, 475–480.

Campeau, E., Ruhl, V.E., Rodier, F., Smith, C.L., Rahmberg, B.L., Fuss, J.O., Campisi, J., Yaswen, P., Cooper, P.K., and Kaufman, P.D. (2009). A versatile viral system for expression and depletion of proteins in mammalian cells. PLoS One 4, e6529.

Cheutin, T., O’Donohue, M.-F., Beorchia, A., Klein, C., Kaplan, H., and Ploton, D. (2003). Three-dimensional organization of pKi-67: a comparative fluorescence and electron tomography study using FluoroNanogold. J. Histochem. Cytochem. 51, 1411–1423.

Croft, J.A., Bridger, J.M., Boyle, S., Perry, P., Teague, P., and Bickmore, W.A. (1999). Differences in the localization and morphology of chromosomes in the human nucleus. J. Cell Biol. 145, 1119–1131.

Cuylen, S., Blaukopf, C., Politi, A.Z., Müller-Reichert, T., Neumann, B., Poser, I., Ellenberg, J., Hyman, A.A., and Gerlich, D.W. (2016). Ki-67 acts as a biological surfactant to disperse mitotic chromosomes. Nature 535, 308–312.

Dohke, K., Miyazaki, S., Tanaka, K., Urano, T., Grewal, S.I.S., and Murakami, Y. (2008). Fission yeast chromatin assembly factor 1 assists in the replication-coupled maintenance of heterochromatin. Genes Cells 13, 1027–1043.

Endl, E., and Gerdes, J. (2000). Posttranslational modifications of the Ki-67 protein coincide with two major checkpoints during mitosis. J. Cell. Physiol. 182, 371–380.

Fazzio, T.G., Huff, J.T., and Panning, B. (2008). An RNAi screen of chromatin proteins identifies Tip60-p400 as a regulator of embryonic stem cell identity. Cell 134, 162–174.

Gerdes, J., Schwab, U., Lemke, H., and Stein, H. (1983). Production of a mouse monoclonal antibody reactive with a human nuclear antigen associated with cell proliferation. Int. J. Cancer 31, 13–20.

Gerdes, J., Lemke, H., Baisch, H., Wacker, H.H., Schwab, U., and Stein, H. (1984). Cell cycle analysis of a cell proliferation-associated human nuclear antigen defined by the monoclonal antibody Ki-67. J. Immunol. 133, 1710–1715.

Ginisty, H., Amalric, F., and Bouvet, P. (1998). Nucleolin functions in the first step of ribosomal RNA processing. EMBO J. 17, 1476–1486.

Ginisty, H., Sicard, H., Roger, B., and Bouvet, P. (1999). Structure and functions of nucleolin. J. Cell Sci. 112 (6), 761–772.

Goto, H., Tomono, Y., Ajiro, K., Kosako, H., Fujita, M., Sakurai, M., Okawa, K., Iwamatsu, A., Okigaki, T., Takahashi, T., et al. (1999). Identification of a novel phosphorylation site on histone H3 coupled with mitotic chromosome condensation. J. Biol. Chem. 274, 25543–25549.

Goto, H., Yasui, Y., Nigg, E.A., and Inagaki, M. (2002). Aurora-B phosphorylates Histone H3 at serine28 with regard to the mitotic chromosome condensation. Genes to Cells 7, 11–17.

Van Hooser, A. a, Yuh, P., and Heald, R. (2005). The perichromosomal layer. Chromosoma 114, 377–388.

Houlard, M., Berlivet, S., Probst, A. V., Quivy, J.-P., Héry, P., Almouzni, G., and Gérard, M. (2006). CAF-1 Is Essential for Heterochromatin Organization in Pluripotent Embryonic Cells. PLoS Genet. 2, e181.

Huang, H., Yu, Z., Zhang, S., Liang, X., Chen, J., Li, C., Ma, J., and Jiao, R. (2010). Drosophila CAF-1 regulates HP1-mediated epigenetic silencing and pericentric heterochromatin stability. J. Cell Sci. 123, 2853–2861.

Isola, J., Helin, H., and Kallioniemi, O.P. (1990). Immunoelectron-microscopic localization of a proliferation-associated antigen Ki-67 in MCF-7 cells. Histochem. J. 22, 498–506.

Kametaka, A., Takagi, M., Hayakawa, T., Haraguchi, T., Hiraoka, Y., and Yoneda, Y. (2002). Interaction of the chromatin compaction-inducing domain (LR domain) of Ki-67 antigen with HP1 proteins. Genes to Cells 7, 1231–1242.

Kaufman, P.D., Kobayashi, R., Kessler, N., and Stillman, B. (1995). The p150 and p60 subunits of chromatin assembly factor I: a molecular link between newly synthesized histones and DNA replication. Cell 81, 1105–1114.

Keller, C., and Krude, T. (2000). Requirement of cyclin/Cdk2 and protein phosphatase 1 activity for chromatin assembly factor 1-dependent chromatin assembly during DNA synthesis. J. Biol. Chem. 275, 35512–35521.

van Koningsbruggen, S., Gierlinski, M., Schofield, P., Martin, D., Barton, G.J., Ariyurek, Y., den Dunnen, J.T., and Lamond, A.I. (2010). High-resolution whole-genome sequencing reveals that specific chromatin domains from most human chromosomes associate with nucleoli. Mol. Biol. Cell 21, 3735–3748.

Krude, T. (1995). Chromatin assembly factor 1 (CAF-1) colocalizes with replication foci in HeLa cell nuclei. Exp. Cell Res. 220, 304–311.

Luo, Y., Ren, F., Liu, Y., Shi, Z., Tan, Z., Xiong, H., Dang, Y., and Chen, G. (2015). Clinicopathological and prognostic significance of high Ki-67 labeling index in hepatocellular carcinoma patients: a meta-analysis. Int. J. Clin. Exp. Med. 8, 10235–10247.

Ma, N., Matsunaga, S., Takata, H., Ono-Maniwa, R., Uchiyama, S., and Fukui, K. (2007). Nucleolin functions in nucleolus formation and chromosome congression. J. Cell Sci. 120, 2091–2105.

MacCallum, D.E., and Hall, P.A. (1999). Biochemical characterization of pKi67 with the identification of a mitotic-specific form associated with hyperphosphorylation and altered DNA binding. Exp. Cell Res. 252, 186–198.

du Manoir, S., Guillaud, P., Camus, E., Seigneurin, D., and Brugal, G. (1991). Ki-67 labeling in postmitotic cells defines different Ki-67 pathways within the 2c compartment. Cytometry 12, 455–463.

Marheineke, K., and Krude, T. (1998). Nucleosome assembly activity and intracellular localization of human CAF-1 changes during the cell division cycle. J. Biol. Chem. 273, 15279–15286.

Matheson, T.D., and Kaufman, P.D. (2015). Grabbing the genome by the NADs. Chromosoma 125, 361–371.

Matsumoto-Taniura, N., Pirollet, F., Monroe, R., Gerace, L., and Westendorf, J.M. (1996). Identification of novel M phase phosphoproteins by expression cloning. Mol. Biol. Cell 7, 1455–1469.

Moggs, J.G., Grandi, P., Quivy, J.P., Jónsson, Z.O., Hübscher, U., Becker, P.B., and Almouzni, G. (2000). A CAF-1-PCNA-mediated chromatin assembly pathway triggered by sensing DNA damage. Mol. Cell. Biol. 20, 1206–1218.

Murzina, N., Verreault, a, Laue, E., and Stillman, B. (1999). Heterochromatin dynamics in mouse cells: interaction between chromatin assembly factor 1 and HP1 proteins. Mol. Cell 4, 529–540.

Németh, A., Conesa, A., Santoyo-Lopez, J., Medina, I., Montaner, D., Péterfia, B., Solovei, I., Cremer, T., Dopazo, J., and Längst, G. (2010). Initial genomics of the human nucleolus. PLoS Genet. 6, e1000889.

Padeken, J., and Heun, P. (2014). Nucleolus and nuclear periphery: Velcro for heterochromatin. Curr. Opin. Cell Biol. 28C, 54–60.

Pezzilli, R., Partelli, S., Cannizzaro, R., Pagano, N., Crippa, S., Pagnanelli, M., and Falconi, M. (2016). Ki-67 prognostic and therapeutic decision driven marker for pancreatic neuroendocrine neoplasms (PNENs): A systematic review. Adv. Med. Sci. 61, 147–153.

Politz, J.C.R., Scalzo, D., and Groudine, M. (2016). The redundancy of the mammalian heterochromatic compartment. Curr. Opin. Genet. Dev. 37, 1–8.

Polo, S.E., Theocharis, S.E., Klijanienko, J., Savignoni, A., Asselain, B., Vielh, P., and Almouzni, G. (2004). Chromatin assembly factor-1, a marker of clinical value to distinguish quiescent from proliferating cells. Cancer Res. 64, 2371–2381.

Pyo, J.-S., Kang, G., and Sohn, J.H. (2016). Ki-67 labeling index can be used as a prognostic marker in gastrointestinal stromal tumor: a systematic review and metaanalysis. Int. J. Biol. Markers 31, e204–10.

Quivy, J.-P., Roche, D., Kirschner, D., Tagami, H., Nakatani, Y., and Almouzni, G. (2004). A CAF-1 dependent pool of HP1 during heterochromatin duplication. EMBO J. 23, 3516–3526.

Quivy, J.-P., Gérard, A., Cook, A.J.L., Roche, D., and Almouzni, G. (2008). The HP1-p150/CAF-1 interaction is required for pericentric heterochromatin replication and Sphase progression in mouse cells. Nat. Struct. Mol. Biol. 15, 972–979.

Reese, B.E., Bachman, K.E., Baylin, S.B., and Rountree, M.R. (2003). The methyl-CpG binding protein MBD1 interacts with the p150 subunit of chromatin assembly factor 1. Mol. Cell. Biol. 23, 3226–3236.

Richards-Taylor, S., Ewings, S.M., Jaynes, E., Tilley, C., Ellis, S.G., Armstrong, T., Pearce, N., and Cave, J. (2016). The assessment of Ki-67 as a prognostic marker in neuroendocrine tumours: a systematic review and meta-analysis. J. Clin. Pathol. 69, 612–618.

Rickards, B., Flint, S.J., Cole, M.D., and LeRoy, G. (2007). Nucleolin is required for RNA polymerase I transcription in vivo. Mol. Cell. Biol. 27, 937–948.

Roger, B., Moisand, A., Amalric, F., and Bouvet, P. (2002). Repression of RNA polymerase I transcription by nucleolin is independent of the RNA sequence that is transcribed. J. Biol. Chem. 277, 10209–10219.

Rolef Ben-Shahar, T., Castillo, A.G., Osborne, M.J., Borden, K.L.B., Kornblatt, J., and Verreault, A. (2009). Two fundamentally distinct PCNA interaction peptides contribute to chromatin assembly factor 1 function. Mol. Cell. Biol. 29, 6353–6365.

Saiwaki, T., Kotera, I., Sasaki, M., Takagi, M., and Yoneda, Y. (2005). In vivo dynamics and kinetics of pKi-67: transition from a mobile to an immobile form at the onset of anaphase. Exp. Cell Res. 308, 123–134.

Sarraf, S. a, and Stancheva, I. (2004). Methyl-CpG binding protein MBD1 couples histone H3 methylation at lysine 9 by SETDB1 to DNA replication and chromatin assembly. Mol. Cell 15, 595–605.

Sheval, E. V, and Polyakov, V.Y. (2008). The peripheral chromosome scaffold, a novel structural component of mitotic chromosomes. Cell Biol. Int. 32, 708–712.

Smith, S., and Stillman, B. (1989). Purification and characterization of CAF-I, a human cell factor required for chromatin assembly during DNA replication in vitro. Cell 58, 15–25.

Smith, S., and Stillman, B. (1991). Immunological characterization of chromatin assembly factor I, a human cell factor required for chromatin assembly during DNA replication in vitro. J. Biol. Chem. 266, 12041–12047.

Smith, C.L., Matheson, T.D., Trombly, D.J., Sun, X., Campeau, E., Han, X., Yates, J.R., and Kaufman, P.D. (2014). A separable domain of the p150 subunit of human chromatin assembly factor-1 promotes protein and chromosome associations with nucleoli. Mol. Biol. Cell 25, 2866–2881.

Sobecki, M., Mrouj, K., Camasses, A., Parisis, N., Nicolas, E., Llères, D., Gerbe, F., Prieto, S., Krasinska, L., David, A., et al. (2016). The cell proliferation antigen Ki-67 organises heterochromatin. Elife 5, e13722.

Sun, H., and Hunter, T. (2012). Poly-small ubiquitin-like modifier (PolySUMO)-binding proteins identified through a string search. J. Biol. Chem. 287, 42071–42083.

Takagi, M., Matsuoka, Y., Kurihara, T., and Yoneda, Y. (1999). Chmadrin: a novel Ki-67 antigen-related perichromosomal protein possibly implicated in higher order chromatin structure. J. Cell Sci. 112 (1), 2463–2472.

Takagi, M., Nishiyama, Y., Taguchi, A., and Imamoto, N. (2014). Ki67 antigen contributes to the timely accumulation of protein phosphatase 1γ on anaphase chromosomes. J. Biol. Chem. 289, 22877–22887.

Takami, Y., Ono, T., Fukagawa, T., Shibahara, K., and Nakayama, T. (2007). Essential role of chromatin assembly factor-1-mediated rapid nucleosome assembly for DNA replication and cell division in vertebrate cells. Mol. Biol. Cell 18, 129–141.

Traut, W., Endl, E., Scholzen, T., Gerdes, J., and Winking, H. (2002). The temporal and spatial distribution of the proliferation associated Ki-67 protein during female and male meiosis. Chromosoma 111, 156–164.

Ugrinova, I., Monier, K., Ivaldi, C., Thiry, M., Storck, S., Mongelard, F., and Bouvet, P. (2007). Inactivation of nucleolin leads to nucleolar disruption, cell cycle arrest and defects in centrosome duplication. BMC Mol. Biol. 8, 66.

Uwada, J., Tanaka, N., Yamaguchi, Y., Uchimura, Y., Shibahara, K., Nakao, M., and Saitoh, H. (2010). The p150 subunit of CAF-1 causes association of SUMO2/3 with the DNA replication foci. Biochem. Biophys. Res. Commun. 391, 407–413.

Verheijen, R., Kuijpers, H.J., Schlingemann, R.O., Boehmer, A.L., van Driel, R., Brakenhoff, G.J., and Ramaekers, F.C. (1989a). Ki-67 detects a nuclear matrix-associated proliferation-related antigen. I. Intracellular localization during interphase. J. Cell Sci. 92 (1), 123–130.

Verheijen, R., Kuijpers, H.J., van Driel, R., Beck, J.L., van Dierendonck, J.H., Brakenhoff, G.J., and Ramaekers, F.C. (1989b). Ki-67 detects a nuclear matrix-associated proliferation-related antigen. II. Localization in mitotic cells and association with chromosomes. J. Cell Sci. 92 (4), 531–540.

